# Characterizing the regulatory logic of transcriptional control at the DNA sequence level by ensembles of thermodynamic models

**DOI:** 10.1101/2025.02.26.640137

**Authors:** Alan Utsuni Sabino, Drielly de Moraes Guerreiro, Ah-Ram Kim, Alexandre Ferreira Ramos, John Reinitz

## Abstract

Understanding how the genome encodes the regulatory logics of transcription is a main challenge of the post-genomic era to be overcome with the aid of customized computational tools. We report an automatic framework for analyzing an ensemble of fittings to data of a thermodynamics-based sequence-level model for transcriptional regulation. The fittings are clustered accordingly with their intrinsic regulatory logics. A multiscale analysis enables visualization of quantitative features resulting from the deconvolution of the regulatory profile provided by multiple transcription factors interacting with the locus of a gene. Quantitative experimental data on reporters driven by the whole locus of the even-skipped gene in blastoderm of Drosophila embryos was used for validating our approach. A few clusters of highly active DNA binding sites within the enhancers collectively modulate even-skipped gene transcription. Analysis of variable enhancers’ length shows the importance of bound protein-protein interactions for transcriptional regulation.

## 1 Introduction

The interpretation of models of natural phenomena containing large numbers of parameters and multiple scales is a central issue in 21st– century science. Although correct predictions by such models are obviously of great utility, their central purpose is to increase human understanding of complex natural phenomena. This issue is particularly acute in systems biology, where a key problem is the elucidation of the control of a variety of processes by DNA sequences, notably including transcription and development. Elegant fundamental studies of developmental processes by parameter free models have been performed (Bauer et al., 2021; McGough et al., 2024; Bialek, 2024; Sokolowski et al., 2025), and raised the existence of quantitative limits to be satisfied by the molecular machinery governing the synthesis of gene products in developing *Drosophila* embryos. However, as this approach cannot address the inherently parametric effects of DNA sequence, it does not shed light on how the astonishing optimality observed in biological processes is reached. Indeed, this is also an issue of models exclusively relying on machine learning techniques, which do not address the specific application domain internal structure of natural science models, and are hard to interpret even in the presence of effective prediction (Liu et al., 2020). Therefore, constructing phenomenological models aiming at a description of the processes underlying regulation of expression in metazoans is a main scientific challenge of the post-genomic era.

Typically, models of complex phenomena with many parameters are constrained by—fit—to observed quantitative data, which itself is subject to experimental uncertainty. Ideally, one aims to obtain a sufficiently large number of fittings for the parameters of the model constrained by data. That essentially comprises a posterior distribution, a picture that is true whether or not the fittings are obtained by an explicit Bayesian method. In simple problems, standard statistical methods can be used. In more complex situations with many parameters, one is typically interested in what classes of fittings are compatible with the data. For that, machine (or statistical) learning techniques may be useful for dimensionality reduction and identification of fittings of a given class. Whether assessed under the light of the phenomenology of a model, those classes enable one to build a scientific interpretation about the mechanisms underpinning complex phenomena. Hence, developing automatic procedures to improve the identification of fittings coherent with the experiments is a key goal of systems biology because its subjects have many parameters and require ensembles of fittings of phenomenological models to be understood.

Here we present a computational framework for the analysis of a multi-parameter and multiscale phenomenon. That is explored in the context of an automated machine interpretation of the genomic regulatory mechanisms as obtained from ensembles of fits to sequence level models of the modulation of gene transcription. In this approach a diffusion limited Arhenius law is employed to model promoter activity mediated by the action of Transcription Factors (TFs) responsible for reducing the energy barrier of transcriptional activation. As inputs, these models receives DNA sequence and the concentration of TFs, along with their Positional Weight Matrices (PWMs) (Zhao et al., 2009), and produce the concentration of mRNAs as output (Reinitz et al., 2003). Such models have been applied to investigate regulation of gene expression in early *Drosophila* embryos (Janssens et al., 2006; Kim et al., 2013; Martinez et al., 2014; Barr and Reinitz, 2017; Barr et al., 2017; Liu et al., 2020; Masuda et al., 2024), Murine erythropoiesis (Bertolino et al., 2016), and human cells (Kang and Kim, 2024). Here we apply our framework using the DNA sequence of the whole locus of the *even-skipped* gene (*eve*) in the blastoderm of the *Drosophila* embryo to fit experimentally determined data on mRNA levels along the anterior-posterior axis (A–P) from stripe 2 to 7 (Barr and Reinitz, 2017). Previous applications reported analysis of few model fits, each involving 32 parameters, and human inspection was employed for determining the compatibility of genomic regulatory activity with experimental data at an enhancer-level resolution (Barr and Reinitz, 2017). Similarly, few fittings for a sector of *eve* locus were found with genomic resolution down to individual binding sites (Kim et al., 2013). However, an automated unified framework to provide an ensemble of experimentally compatible fittings for the *eve* locus DNA sequence at genomic resolution down to binding sites is missed.

In this manuscript, we present an automated toolset to characterize the biological properties of an ensemble of up to several thousand fits that produce output within the uncertainty of the expression data. The biological characterization provided by our method extends to the level of single binding sites, their occupancy by TFs and corresponding physiological consequences. We describe the mathematical formulation of the sequence level thermodynamic model for transcriptional regulation in the Supplementary Information (SI), and its biological meaning in Section 2. The automated method is described in Section 3. In Section 4, we present the biological insights propitiated by our model.

## 2 System and methods

In the SI, we describe the equations comprising our physiological model of transcription. It is based on a thermodynamic picture of the binding of TFs to DNA together with phenomenological rules for the biological function of TFs once bound. The former component gives this class of models its name (Barr and Reinitz, 2017; Barr et al., 2017; Bertolino et al., 2016; Fakhouri et al., 2010; He et al., 2010; Janssens et al., 2006; Kazemian et al., 2010; Kim et al., 2013; Martinez et al., 2014; Reinitz et al., 2003; Samee and Sinha, 2014; Sayal et al., 2016; Segal et al., 2008), while the latter class expresses well known experimental results in computable equations. We next describe how we fit the model to data, and lastly explain how the structure of the model reveals a cluster structure on the actions of TFs in specific instances of fit to data. This information aids one to understand the results presented in Sections 3 and 4. Here we provide a biological description of the inputs, outputs, and layers of information processing contained in this physiological model for transcriptional regulation.

Figure 1 summarizes the flow of our system and the biological implications built upon on its outcomes. The whole sequence of the locus of *eve* gene must be taken into account along with information about the openness of its chromatin, as obtained from two experimental datasets (Li et al., 2011; McKay and Lieb, 2013; Barr and Reinitz, 2017). The regulatory profile is further characterized by determining the minimal set of TFs driving a specific spatio-temporal pattern of transcription. Namely, the following TFs Bicoid (Bcd), Caudal (Cad), *Drosophila*-STAT (Dst), Hunchback (Hb), Giant (Gt), Kruppel (Kr), Knirps (Kni), Tailless (Tll) are used to model the transcriptional regulation of *eve*. The former three TFs are activators, the latter four repressors, and Hb plays a role of repressor or activator if Bcd or Cad are bound to sufficiently close binding sites.

**Fig. 1:**
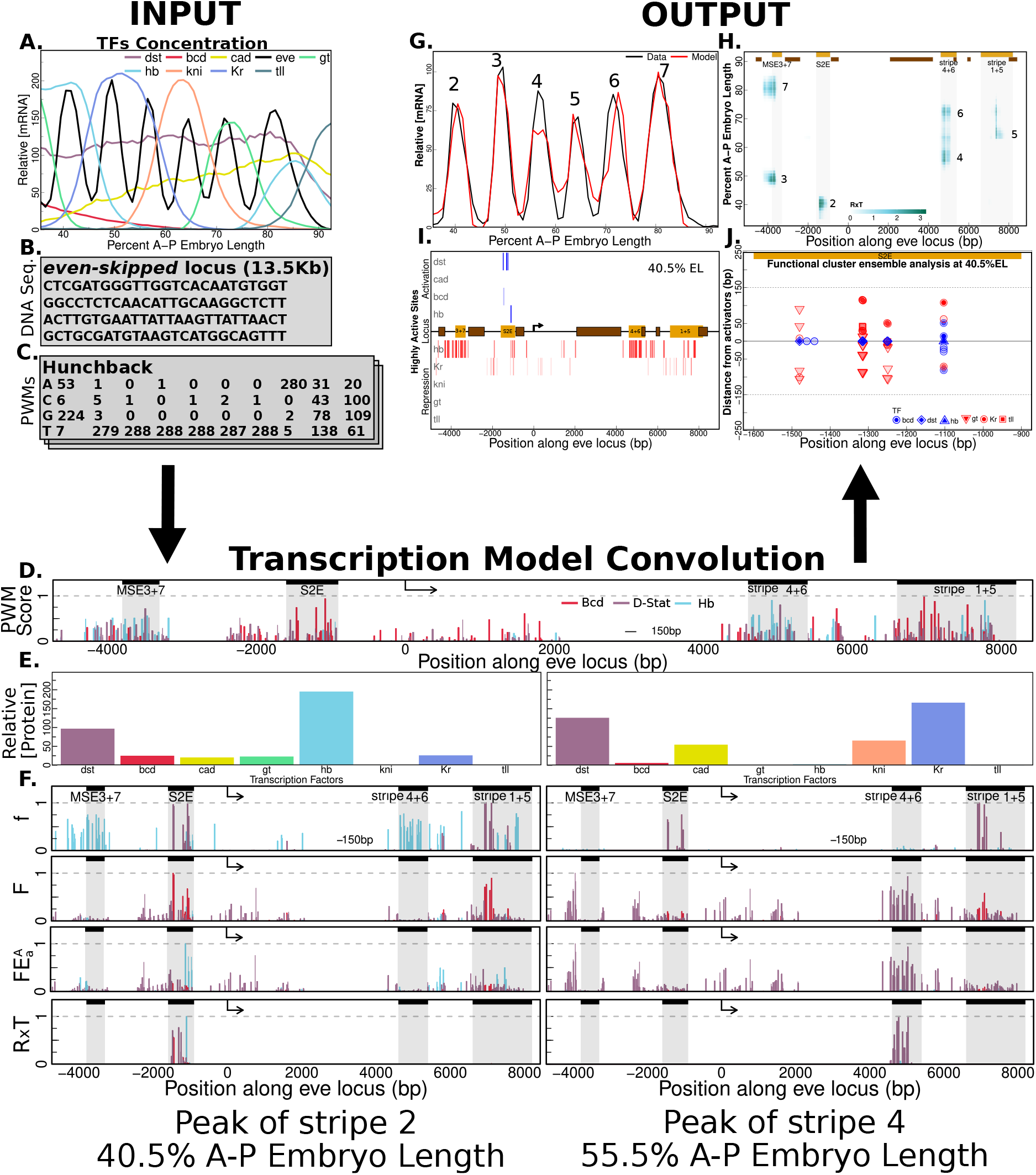
Transcription model convolution scheme. TFs concentration, DNA sequence, and, PWMs are the model’s inputs (A-C). Our model has multiple layers that encode the phenomenology of transcription regulation in fruit fly embryos. (D) We represent the first layer with Bcd, Dst, and, Hb normalized PWM scores. (E) shows protein concentration in two different EL positions along the A–P axis. (F) represents the binding sites activities across the layers at the EL positions corresponding with the protein concentration in (D).

The *inputs* of our system are the protein concentrations of the interacting TFs along A–P (Figure 1**A**), the DNA sequence of the locus of *eve* gene, with closed chromatin regions explicitly indicated (Figure 1**B**), and PWMs corresponding to TFs (Figure 1**C**). The values of the parameters of the model are obtained by application of the Simulated Annealing (SA) algorithm (Kirkpatrick et al., 1983; Chu et al., 1999; Lou and Reinitz, 2016).

The *processing* of the input corresponds to a deconvolution in which the scores of each binding site along the locus, as estimated from the PWMs of TFs along with their concentrations, are governing the activation of the promoter site by specific DNA regions driving transcription along the A–P axis. We exemplify the role of the first layer of regulation, showing the scores of binding sites for Bcd (red), Dst (purple), and Hb (light blue) along the whole locus (see Figure 1**D**). This layer of the model does not encode spatial information *per se*, that is, the scores are the same for all nuclei in the embryo. Positional information depends on the variation of the protein concentrations of the TFs, as we exemplify in Figure 1**E**, for positions 40.5% and 55.5% of the embryo length (EL) along the A–P axis. Hence, understanding the differential activation of the promoter requires integrating sequence-level information with that of available TFs through hierarchically interacting layers as shown in Figure 1**F**. The second layer captures the positional information by means of the occupancy of the binding sites of each TF as shown in the graph for *f* . The next three layers phenomenologically quantify the effects of the interactions among bound TFs as described in SI: coactivation and quenching as encoded by *F* – SI Eq.(4); activation efficiency denoted by 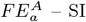 Eq.(5); and enhancer competition indicated by *R* × *T* – SI Eq.(9). In each layer, the relative heights of the vertical bars are dependent of position. In a given position, the relative heights of the bars change as we go through the model layers, and that reveals the importance of composition between the binding sites and their bound TFs on driving transcription of *eve*. Here on that composite will be simply referred to as binding site. Application of the latest layer refines activation regions toward those previously established as enhancers.

Because we employ a sequence level thermodynamic model, the *output* enables a multi-resolution analysis of the mechanisms underlying *eve* transcriptional regulation. In a previous study, the analysis of the spatial pattern of concentration of *eve* and the respective DNA regions driving transcription was presented (Barr and Reinitz, 2017). Figure 1**G** shows that the transcriptional model may recover the experimentally observed spatial patterns of concentration of *eve* along the A–P axis. Position along *eve* locus is shown in the horizontal axis of Figure 1**H**, while the vertical axis indicates position along A–P. The respective DNA regions activating *eve* promoter to express stripes 2 to 7 are highlighted accordingly with their contribution measured as *R* × *T* . Here, we introduce an analysis of the binding sites configurations driving expression in a given spatial position. Figure 1**I** shows the highly active sites contributing to activation of transcription at the peak of stripe 2. The activators are shown in blue and repressors in red. Those *major activators* of transcription are distinguished by the transparency of the vertical bars, and they concentrate within the Stripe 2 Enhancer (S2E) region. DNA regions beyond S2E are densely occupied by Hb and Kr and prevent, mostly by quenching, the action of activators eventually bound there. A zoom in the neighborhood of the binding sites for activators is shown in Figure 1**J**. The vertical axis gives the distance of a neighboring binding site for a quencher or coactivator to the binding site for a major activator. The position along the locus of the site for a major activator is shown in the horizontal axis. Each vertical set of binding sites having a major activating TF at the center along with its quenchers and coactivators above and below the dashed line form a functional cluster. In Section 4 we will discuss the results presented in the output.

## 3 Algorithm and Implementation

Our goal is to obtain an ensemble of parameter values for the model presented in SI to generate results compatible with experimental observations. Those are denoted as *biologically compatible fittings*. Nuclear resolution data on the spatial pattern of expression of *eve* and the TFs modulating its promoter activity were used (Poustelnikova et al., 2005; Surkova et al., 2008a,b; Pisarev et al., 2008) along with experimentally determined binding sites for activators driving the formation of *eve* stripe 2 (Small et al., 1992; Ludwig and Kreitman, 1995). The automated selection of biologically compatible fittings is performed in two rounds of a couple of thousands of applications of SA for parameter optimization (Subsec. 3.1). The set of fittings outcoming from each round of SA is subjected to two filters (Subsec. 3.2). The first filter seeks for appropriate spatial patterns of *eve* expression along the A–P axis, and the second verifies if the activating binding sites are located within the boundaries of experimentally determined enhancers driving expression of stripe 2 (Stanojevic et al., 1991; Small et al., 1992; Ludwig and Kreitman, 1995). That procedure produces an ensemble of biologically compatible fittings which, after being statistically analyzed, enable the reduction of the search regions within the parameter space of the model (Subsec. 3.3). A binding site popularity ranking analysis is then applied to all biologically compatible fittings to investigate both the controlling DNA regions, *i*.*e*., those sequences driving promoter activity, and their respective binding sites in each position along A–P axis. A complete characterization of the sets of interacting binding sites responsible for the formation of *eve* stripes, denoted as *functional clusters*, is automatically provided (Subsec. 3.4). The ensemble of fittings enables the investigation of multiple regulatory logics producing the same expression pattern. That can be done by analyzing the clusterization of biologically compatible fittings within the parameter space (Subsec 3.5). Implementation details are provided in Subsec. 3.6.

## 3.1 Generating an ensemble of fittings

Experimental data about *eve* transcription levels and concentration of its respective TFs were obtained from FlyEx database (Poustelnikova et al., 2005; Surkova et al., 2008a,b; Pisarev et al., 2008). The PWMs for the TFs along with the locus sequence were discussed in Barr and Reinitz (2017). The parameter values for the transcription model presented in SI were obtained by optimization (Sup. Figure 1). We employed the SA technique as in previous studies aiming at investigating regulation of gene expression in *Drosophila* embryos (Jaeger et al., 2004; Janssens et al., 2006; Manu et al., 2009a,b; Kim et al., 2013; Barr and Reinitz, 2017). The SA algorithm aims to minimize Σ*ν* [𝒪_*ν*_ − ℳ _*ν*_]^2^, where 𝒪_*ν*_ and ℳ_*ν*_ respectively indicate relative *eve* mRNA concentrations as experimentally observed and predicted by the model and *?* denotes the percentage of EL along the A–P axis and runs from 35.5 to 92.5% EL. The SA search region within the parameter space was set based on *a priori* biological information (Barr and Reinitz, 2017).

Because the SA is governed by a stochastic process, its output has an intrinsic randomness. Hence, an ensemble of fittings to data can be produced to be statistically analyzed with the aim of hypothesizing the regulatory logics underlying a given spatial pattern of gene transcription. Two issues must be approached: *1*. Not all fittings are biologically compatible, and they must be filtered; *2*. Most promising fittings may be clustered within the parameter space to determine searching regions having a higher probability of generating biologically compatible fittings. Previously, issue *1* was approached manually and based on the analysis of few good fittings to data. As a consequence, issue *2* has been neglected. Here, we approach both issues by means of an automatic procedure.

### 3.2 Automatic selection of biologically compatible fittings

The biological compatibility of the fittings obtained by SA is determined by the application of two filters. The first filter selects candidate fittings based on the Root Mean Square (RMS) value comparing theoretical and experimentally determined *eve* stripe patterns of transcription. That number enables the selection of fittings generating spatial patterns within experimental error.

The fittings selected in the previous step will pass through the second filter because the controlling DNA regions being activated may not correspond to those experimentally known (Small et al., 1992; Ludwig and Kreitman, 1995). Hence, for a given fitting, the program lists the highly active binding sites contributing to transcriptional activity, here set as those causing 80% of the total reduction for transcription initiation along A–P axis. That percentage is a free parameter of our framework. Only fittings whose transcriptional activity at the peak of stripe 2 had a binding site located within the S2E region among the major activators are held as biologically compatible.

### 3.3 Biologically guided reduction of the parameter space volume

We performed two rounds of 2000 fittings to data by SA. The first round produced 81 biologically compatible fittings. The parameter values of biologically compatible fittings are distributed either *a)* having most of its values lying within some interval while presenting few outliers; or *b)* not having outliers. For the parameters belonging to category *a)* we set two reference values Max = *Q*_3_ + 1.5 IQ and Min = *Q*_1_ − 1.5 IQ, and redefined the limits of search range in terms of Max and/or Min, where *Q*_1_ (*Q*_3_), is the first (third) quartile and *IQ* = *Q*_3_ − *Q*_1_. For the parameters belonging to category *b)* we used the largest and/or smallest estimated parameter values for defining their search range. The new search ranges were used in the second round of annealings, and we produced 1881 biologically compatible fittings. In the first round we produced 81, which is a 23–fold change after search region refinement.

### 3.4 Ensemble analysis of biologically compatible models

A binding site popularity ranking analysis of the biologically compatible fittings is performed for discovering the logics of regulation of transcription of *eve* gene along the A–P axis. The highly active binding sites driving expression in each position of the A–P axis are collected from each fitting. Then, a weighted sum of the contribution of each activating binding site is performed. The weight of a binding site is its contribution to energy barrier reduction in each fitting, multiplied by the number of times of appearance of this binding site in all fittings. The resulting number is divided by the total number of occurrences of the binding site that appears the most among all biologically compatible fittings.

The above procedure produces the set of most popular highly active binding sites driving transcription in each position along A–P axis. The regulatory logics underpinning stripe formation is proposed after identification of all binding sites having bound quenchers and coactivators that interact with a specific activator. The combination of each set composed by a highly active binding site, its quenchers, and coactivators, along with their interactive mechanisms, compose what we denote as a functional cluster.

### 3.5 The regulatory logics are further determined based on the clusters of fittings

The large ensemble of biologically compatible fittings enabled the construction of histograms of parameter values. That shedded light on the multi-modality of some parameter values distributions. That prompted us to investigate the formation of clusters of fittings among the biologically compatible models. This analysis revealed that the formation of clusters reflects on different sets of highly active binding sites driving expression along the A–P axis, and that represents different regulatory logics.

The analysis of the clusterization of biologically compatible fittings was performed using *k*-means. For that, we used a *z*-transformation in each parameter value so that they will all be on the same scale. To define the optimal number of clusters, we vary the number of the *k* parameter in the *k*-means algorithm from 2 to 10 using the package NbClust (Charrad et al., 2014) where multiple metrics to assess the number of optimal clusters were employed. A Principal Component Analysis (PCA) was performed using the parameter values of the biologically compatible fittings. The three first principal components were used for visualization of the data, *i*.*e*., ensembles of parameter values, in a lower dimensional space to enable the inspection of the clusters. Based on frequency analysis of the best *k* estimated with NbClust and inspection of PCA clusters, the number of clusters to be set in *k*-means was defined as 3.

### 3.6 Implementation

SA was implemented previously in C++ using the Lam-Delosme algorithm (Lam and Delosme, 1988a,b; Chu et al., 1999; Lou and Reinitz, 2016). The transcription model implementation used as the base version for this work was reported in Barr and Reinitz (2017) and written in C++ with an R interface. We created an environment using Docker to couple these previously developed programs with all necessary packages to run SA, the transcription model, and the R interface.

Our automatic data processing algorithm was implemented as R scripts. All required packages for our analysis were included in a Docker environment. Our algorithm performed all calculations in serial. The runs were performed in batches using our cluster of five machines (see SI for system specification).

## 4 Discussion

### Automatic ensemble analysis shows that model fittings recovering enhancer regions at *eve* locus are in a subdomain of the search space

Figure 2 presents a scheme of a new automatic method to generate biologically compatible fittings for the TM and illustrates its usefulness by the analysis of data on transcription driven by *eve* whole locus. We propose a two-step approach: *i*. to generate an initial set of biologically compatible fittings, and *ii*. to build the distributions of the parameter values to reduce the search space for a second application of our algorithm for finding biologically compatible fittings. Figure 2A shows the inputs of our framework.

**Fig. 2:**
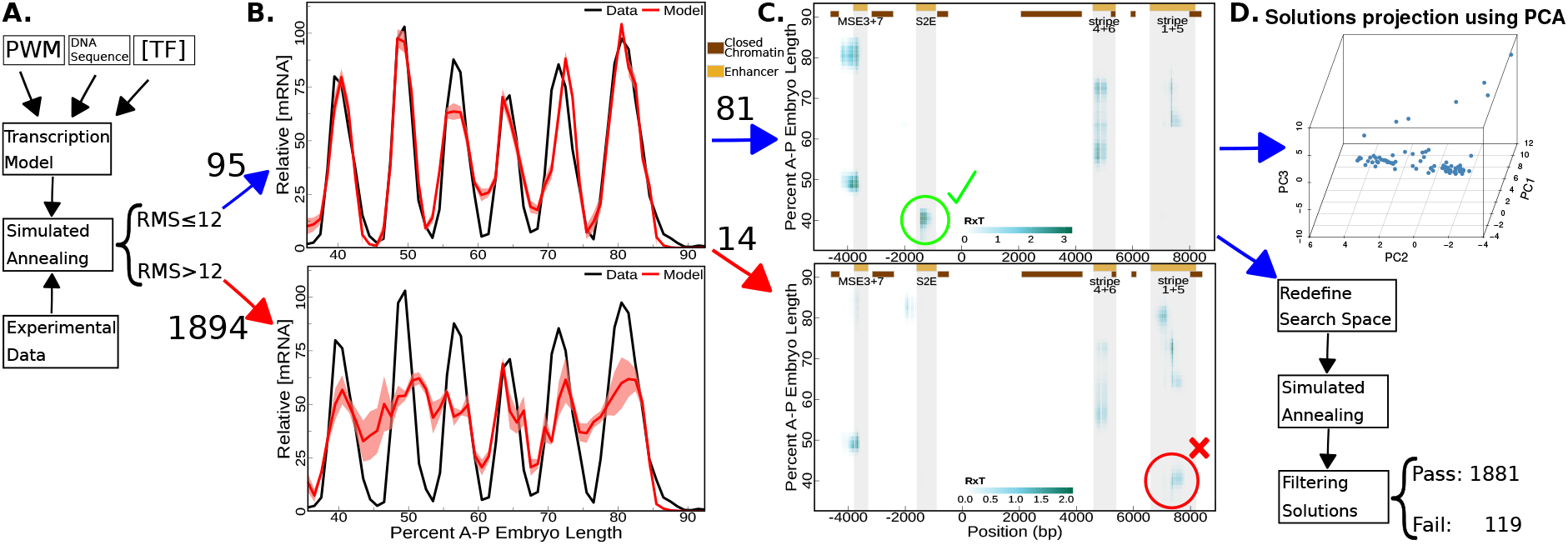
Flow of analysis for obtaining biologically compatible fittings. The flowchart presents our computerized method for automatic filtering and optimization of fittings for the transcriptional model. (A) Simplified scheme of input data and computational process. (B) Mean (red) ± standard deviation (shaded red) of predicted *eve* expression along EL by model fittings with low (top) and high (bottom) RMS values – first filter. Experimental data (black) is shown as reference. (C) Prediction of active regions along *eve* locus. The top (bottom) graphs are highlighting regions of binding sites activation that are compatible (incompatible) with experiments – second filter. (D) 3D Projection of parameter values along first three principal components obtained after application of PCA to biologically compatible fittings (top). Scheme of the process of search space reduction and new round of SA (bottom).

The TM has 32 free parameters to be optimized by SA for fitting the spatial pattern of *eve* stripes (see black lines in Figure 2B). The RMS threshold was set as 12 based on inspection of the graphs (see Figure 2B top). Only 4.78% (95) of the fittings crossed the established threshold used as a first filter. Mean fit predictions (red), with standard deviation, and experimental data (black) are presented in Figure 2B: the top graph shows fittings with good expression prediction (RMS ≤ 12), while representative bad predictions (RMS ≥ 20) are shown at the bottom.

*In silico* sequence-level analysis reveals that some fittings (14.74%) with a good expression pattern do not correctly predict the regions of regulatory activity of *eve* locus DNA sequence corresponding to experimentally established enhancer regions. Figure 2C exhibits classes of samples of fittings which region of the gene locus that is driving the formation of stripe 2 do, or do not, coincide to the enhancer region described in literature. Top graph shows regulatory activity from the stripe 2 enhancer region, indicated by the shaded region along DNA sequence positioned between -1600 and -800 basepairs (bp), at the stripe 2 peak of expression, 40.5% EL along the A–P axis (see Figure 2B top for reference). Stripe 1+5 enhancer has regulatory activity at the peak of expression of stripe 2 in the bottom graph, while stripe 2 enhancer has no regulatory activity. Hence, it shows a class of fittings that are not compatible with the experimental data. After challenging the predicted region regulating the formation of the peak of stripe 2 in comparison with the enhancer region described in literature, just 4.07% (81) fittings produced in the first round of application of SA were selected.

PCA was employed to investigate the biologically compatible fittings in a visualizable space having reduced dimensionality, Figure 2D top. Inspection revealed the existence of clusters of fits to data. After applying the reduction in the search space, 94.05% of model fittings passed the two quality filters.

### Sets of similar fittings to data as determined by clusterization algorithms enabled to hypothesize about different regulatory logic at the locus

In Figure 3, we show the highly activated binding sites along *eve* locus at the peak of stripe 2. Graph **A** shows the binding site popularity ranking analysis of highly active binding sites from the ensemble of fits, while graphs **B–D** indicate the results of applying our algorithm within the sets of similar fittings to data. Our model indicates that Hb, Dst, and Bcd, in decreasing order of contribution, are the main activators of transcription at the peak of stripe 2. Hb is coactivated by Bcd (Kim et al., 2013) (graphs **A, D**). In the ensemble (graph **A**), set 1 (graph **B**), and set 3 (graph **D**) of fits, the highly active sites for activators (blue bars) are concentrated within the region of S2E, while some activation beyond this enhancer is shown in the set 2 (graph **C**). The coincidence of highly active binding sites in graphs **A** and **D** is because set 3 has the largest number of fits. Though there are binding sites for activators along the whole locus, they are not active mainly because of quenching. Hb is the major responsible for quenching transcriptional activity being induced by non-S2E enhancers. Kr is also important because of its higher affinity to binding sites within the region of the enhancers 1 + 5 and S2E.

**Fig. 3:**
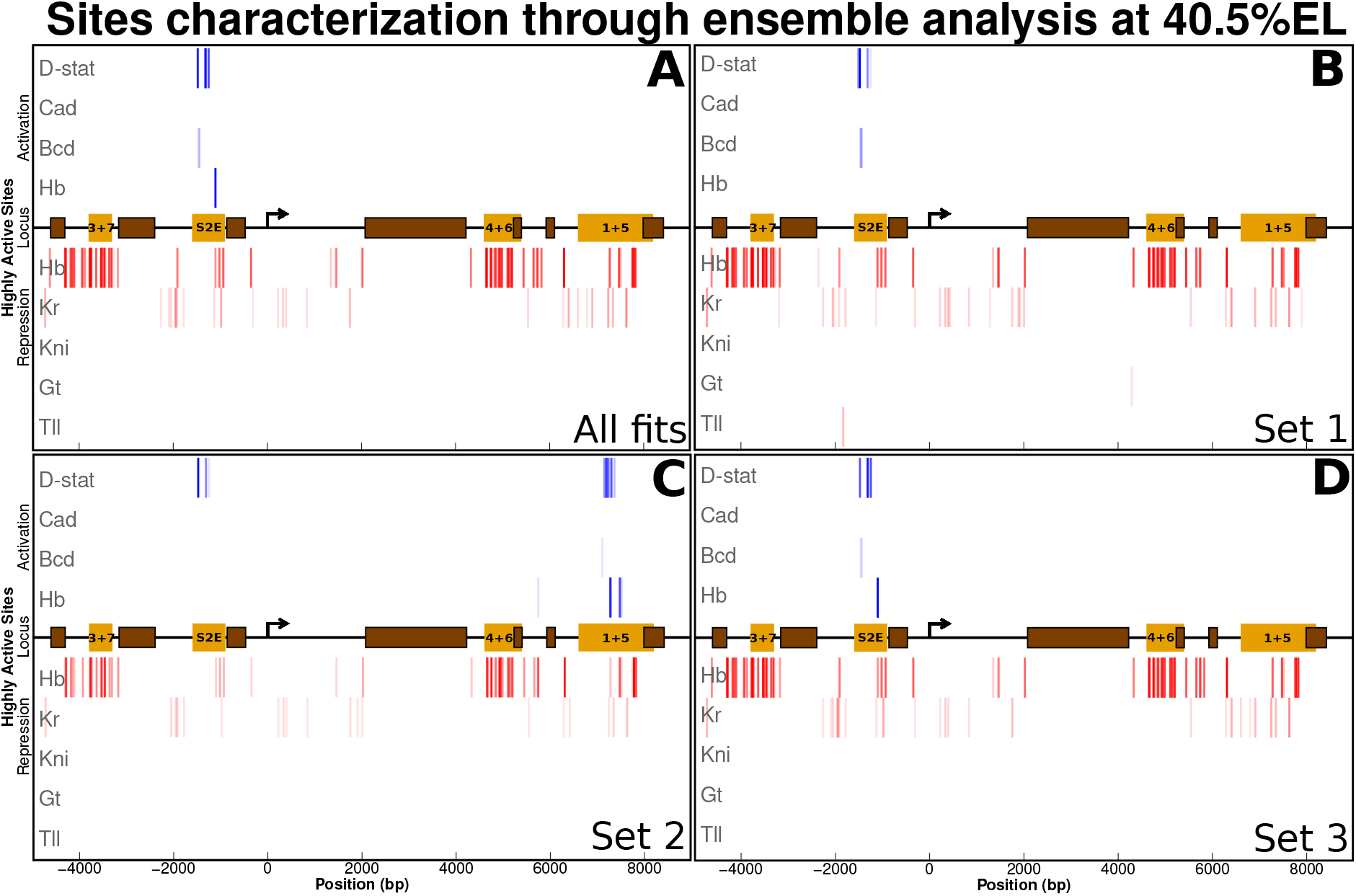
Binding sites interacting to reduce 80% of the energy barrier of transcription activation. The horizontal black line indicates the *eve* locus, with the yellow and brown rectangles respectively denoting the enhancers and the closed chromatin regions. Each row along the vertical axis denotes a TF. The activators are indicated by blue vertical bars, while the red bars show the sites for repressors. The more solid the blue (red) bar, the higher is its activation (or repression efficiency, 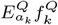). (A–D) shows the ensemble of fittings (A) and its respective sets resulting from clustering parameter values of biologically compatible fittings (B–D).

In Figure 3, the set 2 of fits has an activation of transcription being induced by the binding sites within the enhancer 1 + 5. The top contributors for that activation are Dst and Hb, the latter being coactivated by Bcd (see SI). This set of fits would be discarded for further analysis of the logic of transcriptional regulation of *eve*. But that would be an underestimation of the role of the sets of potentially useful fittings if one is willing to employ our methodology to the study of a new system. The clusterization enables one to collect the fits by their similarity. That can be useful in experiments that investigate the logics underpinning modulation of gene expression by multiple TFs.

Previous experiments have established that the formation of *eve* stripes is driven by enhancers being about ∼ 700 bp long (Stanojevic et al., 1991; Small et al., 1992). The results shown in Figure 3, however, indicate that only a few activating binding sites are the major contributors to *eve* transcription at the peak of stripe 2, for example. Next, we identify functional clusters encompassing collections of binding sites for repressors, activators, and coactivators whose combined effect underlies the modulation of gene transcription by a controlling DNA region.

### Functional cluster characterization aids in understanding the distinctive logics of transcriptional regulation of the locus

In Figure 4, we show the functional clusters that are responsible for an 80% reduction in the energy barrier for transcriptional activation at the peak of stripe 2. Graph **A** shows the binding site popularity ranking analysis of highly active binding sites from the ensemble of fits, while graphs **B–D** indicate the outcomes of application of our algorithm to the sets 1–3 of fittings to data. In each graph, the functional clusters are labeled accordingly with their energy barrier reduction ranking. The activator in a functional cluster is set as the referential blue symbol in the black horizontal line in the middle of the graphs. In general, the functional clusters are composed by a single activator being surrounded by its respective quenchers. However, since Hb is coactivated by Bcd its corresponding functional clusters in graphs 1A and 1D are composed by quenching and activating binding sites. Dst and Bcd binding sites are only surrounded by quenchers, and because we show the stronger repressors, some of their sites are alone. Again, since the set 3 of fits has a larger number of components, it has a stronger influence on determining the functional clusters as obtained from all fits (see graphs **A** and **D**). In both sets 1 and 2, Hb functional clusters do not appear as the highly active sites within the region of S2E. Note that some Hb functional clusters were highly active in the region of the enhancer 1 + 5 (see Figure 3).

**Fig. 4:**
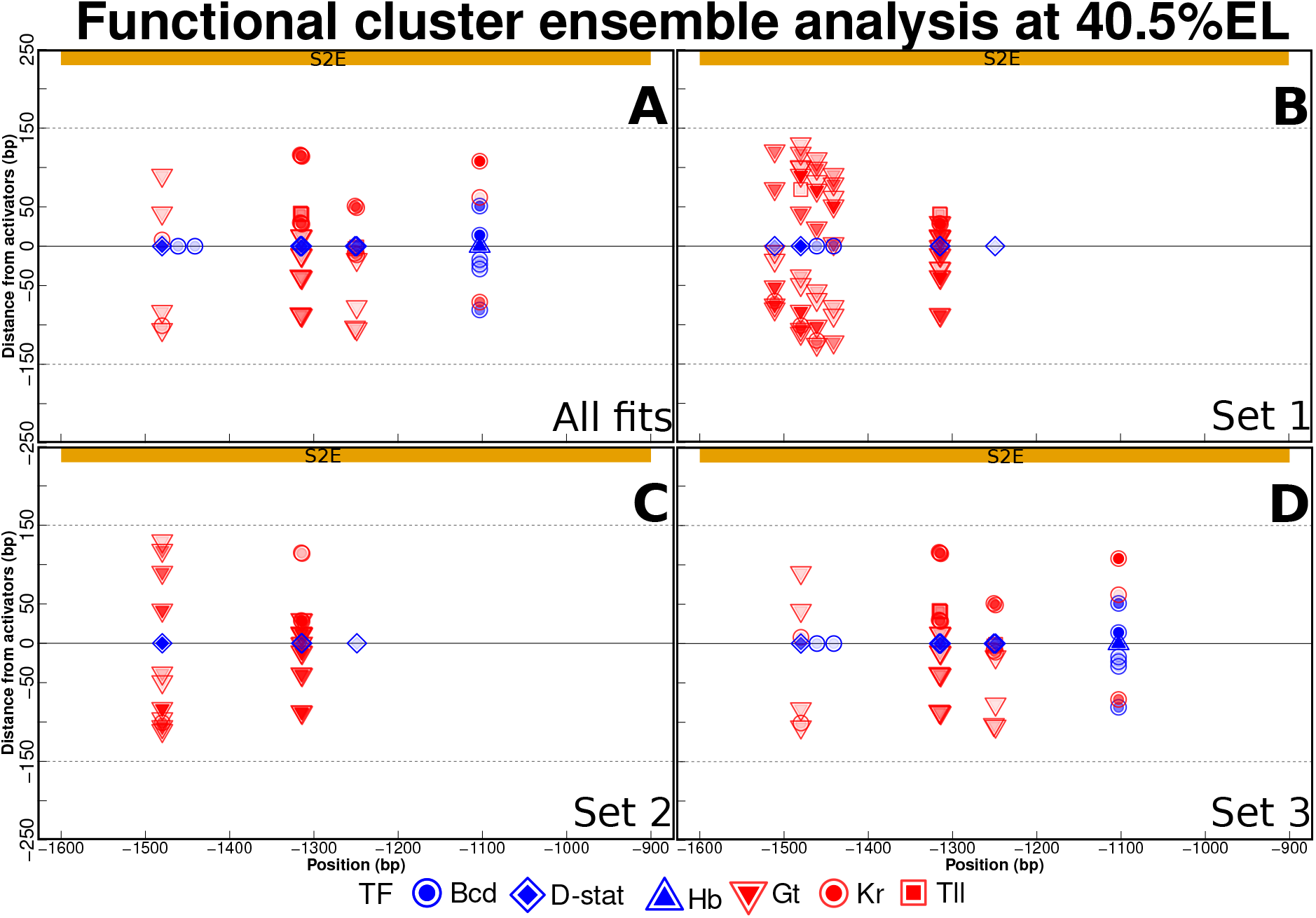
Functional clusters characterization of stripe 2. The top yellow rectangle indicates the region of S2E, and the horizontal axis gives the position along the *eve* locus. The vertical axis indicates the distance, in basepairs, of a binding site to its interacting activator site. The more solid the inner symbol, strength the effect of activation (blue) or repression (red). (A–D) shows fittings ensemble and its sets clustered by parameters.

The functional cluster analysis of Figure 4 enables a higher genomic resolution for understanding the regulatory logic as predicted by the physiological model for regulation of transcription in *Drosophila*. It provides a strategy for selection of the most promising fits to data. For example, graphs **A** and **D** show that the coactivation of **Hb** is a major effect underlying the expression of *eve* at the peak of stripe 2, as previously discussed (Kim et al., 2013). One applying our framework to a new system for investigating the main binding sites driving some gene expression pattern might use this class of graphs for identifying the most promising DNA regions to be experimentally modified. Graphs **B** and **C** show that the spatial patterns of expression of *eve* may be achieved by means of alternative regulatory logics. Under a theoretical perspective, this exemplifies the redundancy of the gene regulatory code and challenges experimentalists to find whether these are model artifacts or observable phenomena.

The functional clusters in Figure 4 may encourage one to investigate whether enhancers of about 700 bp are the outcome of genomic low resolution of experimental techniques. Previous experiments have shown that *eve* stripe two is driven by the S2E region (Ludwig and Kreitman, 1995; Ludwig et al., 1998). The functional clusters are spread over the S2E region, though they may not fulfill it completely. Indeed, experimental results show that a smaller region called MSE2 may drive *eve* stripe 2 (Small et al., 1992), though at lower mRNA levels (Kim, 2012). In the SI, we discussed the window size parameter *α* which indicates the length of DNA segments competing for reducing the energy barrier of promoter activation. Hence, one may vary *α* to investigate the capacity of a given DNA segment to induce transcription. Our framework provides the tools for systematically performing such an analysis, as we show in the next subsection.

### Variation of the length of DNA that influences the promoter shows the importance of bound protein-protein interaction for transcriptional regulation

In Figure 5, we show the effect of changing the parameter *α*. Our model fits *eve* expression while estimating parameter values for each *α*. That leads to distributions of parameter values being dependent on the window size. As *α* increases, more binding sites for activators will be available and, hence, more quenching will be needed to keep the transcriptional activity invariant. Shorter interacting regions (300 bp and 500 bp, Figures 5A and 5B) reveal the major importance of Hb, and hence coactivation, to the formation of the peak of stripe 2. Besides, most of the quenching is performed by Hb with a secondary role being played by Tll. As the window size gets longer (800 bp and 1 kb, Figures 5C and 5D), Dst binding sites start giving a larger contribution for reducing the activation energy barrier, while coactivated Hb action loses importance. Besides, quenching by Hb and Kr from nearby the 5’ border of S2E starts to participate of the functional clusters determining expression at the peak of stripe 2. That is related to the emergence of Dst as a major activator along with its quenchers. Quenching from Hb binding sites are mostly conserved as we increase the window size, while some Kr quenching starts to appear as a compensation for the increased DNA region size inducing promoter activity. This behavior of the model unveils the importance of bound protein-to-protein interactions for transcriptional regulation. It reveals the importance of coactivation when we use smaller window sizes, and how the almost uniform concentration of Dst along the A–P axis raises the necessity of additional quenching when *α* increases. It also shows the limitations of only considering the PWMs as a major guide for establishing regulation of gene expression in metazoans.

**Fig. 5:**
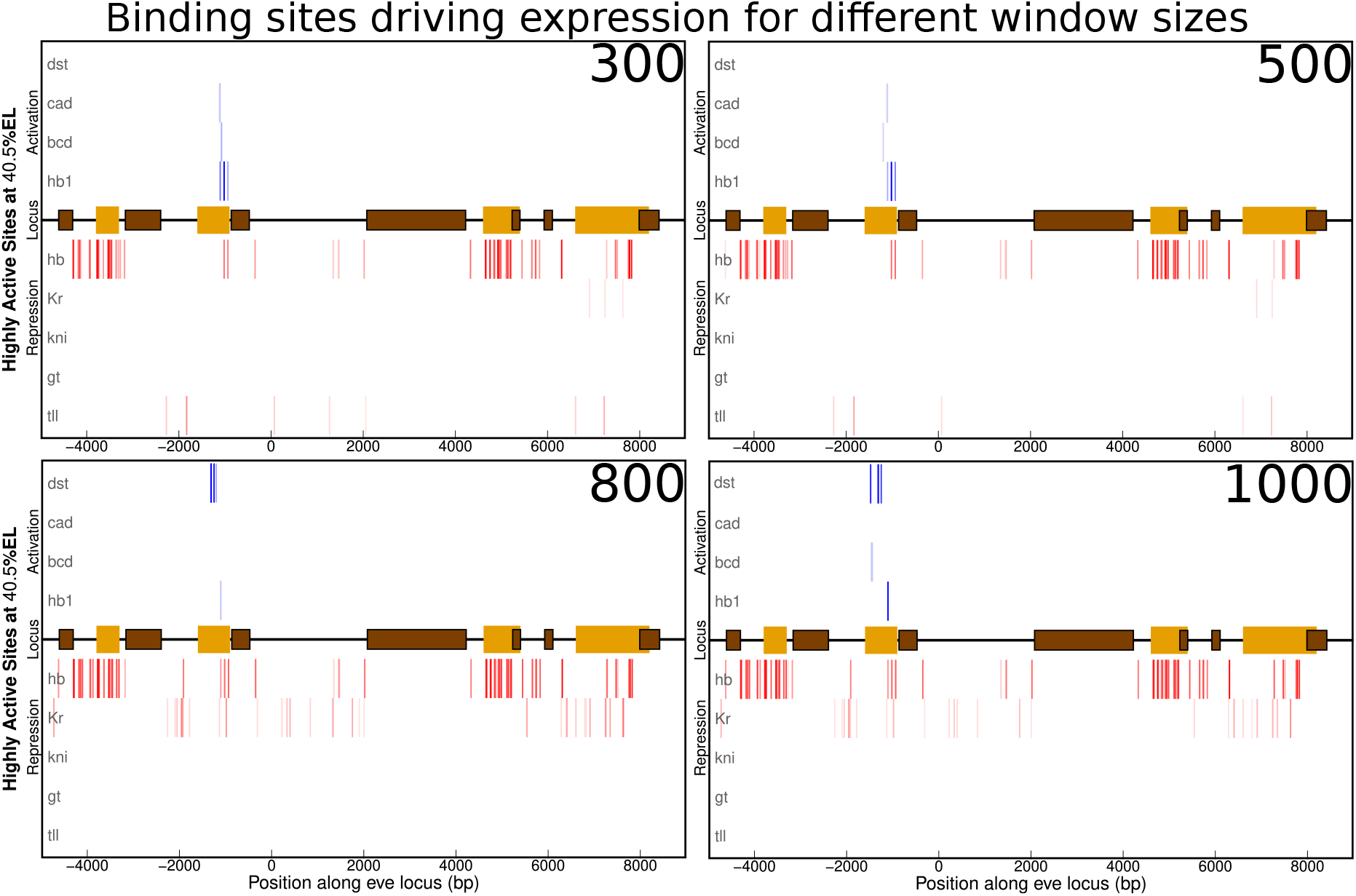
Highly active sites of ensemble fits varying the window size. Binding sites most significant to expression at the peak of stripe 2 are shown for window size parameter with values of 300, 500, 800, and 1000 (A–D).

## Author contributions statement

Methodology, A.U.S., A.F.R. and J.R.; Formal Analysis, A.U.S., A.R.K., A.F.R. and J.R.; Software, D.M.G. and A.U.S.; Writing - Original Draft, A.U.S., A.F.R and J.R.; Writing - Review & Editing, A.U.S. and A.F.R.; Supervision, A.F.R. and J.R.; Project Administration, A.F.R. and J.R.; Funding Acquisition, A.F.R. and J.R.

## Acknowledgments

We would like to thank Kenneth Barr for technical support with newest model implementation as well as Chan-Koo Kang for environment files for configuration of the legacy model implementation, and Luiz P. M. Andrioli for invaluable discussion. This work was supported by funds from the National Institutes of Health R01 OD010936.

## References

K. A. Barr and J. Reinitz. A sequence level model of an intact locus predicts the location and function of nonadditive enhancers. PLoS One, 12:e0180861, 2017. doi:10.1371/journal.pone.0180861. PMCID:PMC5513433.

K. A. Barr, C. Martinez, J. R. Moran, A. R. Kim, A. F. Ramos, and J. Reinitz. Synthetic enhancer design by in silico compensatory evolution reveals flexibility and constraint in cis-regulation. BMC Systems Biology, 11:116, 2017. PMCID:PMC5708098 doi:10.1186/s12918-017-0485-2.

Marianne Bauer, Mariela D. Petkova, Thomas Gregor, Eric F. Wieschaus, and William Bialek. Trading bits in the readout from a genetic network. Proceedings of the National Academy of Sciences, 118:e2109011118, 2021.

E. Bertolino, J. Reinitz, and Manu. The analysis of novel distal Cebpa enhancers and silencers using a transcriptional model reveals the complex regulatory logic of hematopoietic lineage specification. Developmental Biology, 413:128–144, 2016. doi:10.1016/j.ydbio.2016.02.030. PMCID:PMC4878123.

William Bialek. Ambitions for theory in the physics of life. SciPost Physics Lecture Notes, page 84, 2024.

Malika Charrad, Nadia Ghazzali, Véronique Boiteau, and Azam Niknafs. NbClust: An R package for determining the relevant number of clusters in a data set. Journal of Statistical Software, 61:1–36, 2014.

K. W. Chu, Y. Deng, and J. Reinitz. Parallel simulated annealing by mixing of states. The Journal of Computational Physics, 148:646–662, 1999.

Walid D. Fakhouri, Ahmet Ay, Rupindar Sayal, Jacqueline Dresch, Evan Dayringer, and David N. Arnosti. Deciphering a transcriptional regulatory code: modeling short-range repression in the Drosophila embryo. Molecular Systems Biology, 6:341, 2010. PMID:20087339 PMCID:PMC2824527 doi:10.1038/msb.2009.97.

X. He, M. A. H. Samee, C. Blatti, and S. Sinha. Thermodynamics-based models of transcriptional regulation by enhancers: The roles of synergistic activation, cooperative binding and short-range repression. PLoS Computational Biology, 6:e1000935, 2010. PMCID:PMC2940721.

J. Jaeger, S. Surkova, M. Blagov, H. Janssens, D. Kosman, K. N. Kozlov, Manu, E. Myasnikova, C. E. Vanario-Alonso, M. Samsonova, D. H. Sharp, and J. Reinitz. Dynamic control of positional information in the early Drosophila embryo. Nature, 430:368–371, 2004.

H. Janssens, S. Hou, J. Jaeger, A. R. Kim, E. Myasnikova, D. Sharp, and J. Reinitz. Quantitative and predictive model of transcriptional control of the Drosophila melanogaster even skipped gene. Nature Genetics, 38:1159–1165, 2006.

Chan-Koo Kang and Ah-Ram Kim. Deep molecular learning of transcriptional control of a synthetic CRE enhancer and its variants. iScience, 27, 2024.

M. Kazemian, C. Blatti, A. Richards, M. McCutchan, N. Wakabayashi-Ito, A. S. Hammonds, S. E. Celniker, S. Kumar, S. A. Wolfe, M. H. Brodsky, and S. Sinha. Quantitative analysis of the Drosophila segmentation regulatory network using pattern generating potentials. PLoS Biology, 8:e1000456, 2010. PMCID:PMC2923081.

A. R. Kim. Breaking the genomic cis-regulatory code by an experimental and theoretical analysis of eve enhancer fusions. PhD Thesis, Department of Biochemistry and Cell Biology, Stony Brook University, 2012.

A. R. Kim, C. Martinez, J. Ionides, A. F. Ramos, M. Z. Ludwig, N. Ogawa, D. H. Sharp, and J. Reinitz. Rearrangements of 2.5 kilobases of noncoding DNA from the Drosophila even-skipped locus define predictive rules of genomic cis-regulatory logic. PLoS Genetics, 9:e1003243, 2013. PMCID:PMC3585115.

S. Kirkpatrick, C. D. Gelatt, and M. P. Vecchi. Optimization by simulated annealing. Science, 220:671–680, 1983.

J. Lam and J.-M. Delosme. An efficient simulated annealing schedule: Derivation. Technical Report 8816, Yale Electrical Engineering Department, New Haven, CT, September 1988a.

J. Lam and J.-M. Delosme. An efficient simulated annealing schedule: Implementation and evaluation. Technical Report 8817, Yale Electrical Engineering Department, New Haven, CT, September 1988b.

X. Y. Li, S. Thomas, P. J. Sabo, M. B. Eisen, J. A. Stamatoyannopoulos, and M. D. Biggin. The role of chromatin accessibility in directing the widespread, overlapping patterns of Drosophila transcription factor binding. Genome Biology, 12:1–17, 2011. PMID:21473766 PMCID:PMC3218860 doi:10.1186/gb-2011-12-4-r34.

Yi Liu, Kenneth Barr, and John Reinitz. Fully interpretable deep learning model of transcriptional control. Bioinformatics, 36:i499–i507, 2020. doi: 10.1093/bioinformatics/btaa506 PMCID:PMC7355248.

Zhihao Lou and John Reinitz. Parallel simulated annealing using an adaptive resampling interval. Parallel computing, 23:23–31, 2016. doi:10.1016/j.parco.2016.02.001. PMCID:PMC4770898.

M. Z. Ludwig and M. Kreitman. Evolutionary dynamics of the enhancer region of even-skipped in Drosophila. Molecular Biology and Evolution, 12:1002–1011, 1995.

M. Z. Ludwig, N. H. Patel, and M. Kreitman. Functional analysis of eve stripe 2 enhancer evolution in Drosophila: rules governing conservation and change. Development, 125:949–958, 1998.

Manu, S. Surkova, A. V. Spirov, V. Gursky, H. Janssens, A. Kim, O. Radulescu, C. E. Vanario-Alonso, D. H. Sharp, M. Samsonova, and J. Reinitz. Canalization of gene expression in the Drosophila blastoderm by gap gene cross regulation. PLoS Biology, 7:e1000049, 2009a. doi:10.371/journal.pbio.1000049 PMCID:PMC2653557.

Manu, S. Surkova, A. V. Spirov, V. Gursky, H. Janssens, A. Kim, O. Radulescu, C. E. Vanario-Alonso, D. H. Sharp, M. Samsonova, and J. Reinitz. Canalization of gene expression and domain shifts in the Drosophila blastoderm by dynamical attractors. PLoS Computational Biology, 5:e1000303, 2009b. doi:10.1371/journal.pcbi.1000303 PMCID:PMC2646127.

Carlos Martinez, Ah-Ram Kim, Joshua S. Rest, Michael Ludwig, Martin Kreitman, Kevin White, and John Reinitz. Ancestral resurrection of the Drosophila S2E enhancer reveals accessible evolutionary paths through compensatory change. Molecular Biology and Evolution, 31:903–916, 2014. doi: doi:10.1093/molbev/msu042. PMCID:PMC3969564.

Lauro Hiroshi Pimentel Masuda, Alan Utsuni Sabino, John Reinitz, Alexandre Ferreira Ramos, Ariane Machado-Lima, and Luiz Paulo Andrioli. Global repression by tailless during segmentation. Developmental Biology, 505:11–23, 2024.

Lauren McGough, Helena Casademunt, Miloš Nikolić, Zoe Aridor, Mariela D. Petkova, Thomas Gregor, and William Bialek. Finding the last bits of positional information. PRX Life, 2:013016, 2024.

Daniel J. McKay and Jason D. Lieb. A common set of DNA regulatory elements shapes Drosophila appendages. Developmental Cell, 27:306–318, 2013.

A. Pisarev, E. Poustelnikova, M. Samsonova, and J. Reinitz. FlyEx, the quantitative atlas on segmentation gene expression at cellular resolution. Nucleic Acids Research, 37:D560–D566, 2008. PMCID:PMC2686593.

E. Poustelnikova, A. Pisarev, M. Blagov, M. Samsonova, and J. Reinitz. Flyex database. http://urchin.spbcas.ru/flyex, 2005.

J. Reinitz, S. Hou, and D. H. Sharp. Transcriptional control in Drosophila. ComPlexUs, 1:54–64, 2003.

M. A. H. Samee and S. Sinha. Quantitative modeling of a gene’s expression from its intergenic sequence. PLoS Computational Biology, 10:1–21, 2014.

R. Sayal, J. M. Dresch, I. Pushel, B. R. Taylor, and D. Arnosti. Quantitative perturbation-based analysis of gene expression predicts enhancer activity in early Drosophila embryo. eLife, 5:e08445, 2016. PMID:27152947 PMCID:PMC4859806 doi:10.7554/eLife.08445.

E. Segal, T. Raveh-Sadka, M. Schroeder, U. Unnerstall, and U. Gaul. Predicting expression patterns from regulatory sequence in Drosophila segmentation. Nature, 451:535–540, 2008. PMID:18172436 doi:10.1038/nature06496.

S. Small, A. Blair, and M. Levine. Regulation of even-skipped stripe 2 in the Drosophila embryo. The EMBO Journal, 11:4047–4057, 1992.

Thomas R. Sokolowski, Thomas Gregor, William Bialek, and Gašper Tkačik. Deriving a genetic regulatory network from an optimization principle. Proceedings of the National Academy of Sciences, 122:e2402925121, 2025.

D. Stanojevic, S. Small, and M. Levine. Regulation of a segmentation stripe by overlapping activators and repressors in the Drosophila embryo. Science, 254:1385–1387, 1991.

S. Surkova, D. Kosman, K. Kozlov, Manu, E. Myasnikova, A. Samsonova, A. Spirov, C. E. Vanario-Alonso, M. Samsonova, and J. Reinitz. Characterization of the Drosophila segment determination morphome. Developmental Biology, 313(2):844–862, 2008a. PMCID:PMC2254320.

S. Surkova, E. Myasnikova, H. Janssens, K. N. Kozlov, A. Samsonova, J. Reinitz, and M. Samsonova. Pipeline for acquisition of quantitative data on segmentation gene expression from confocal images. Fly, 2:58–66, 2008b. PMCID:PMC2803333.

Y. Zhao, D. Granas, and G. D. Stormo. Inferring binding energies from selected binding sites. PLoS Computational Biology, 5:e1000590, 2009. PMCID:PMC2777355.

