## Supplementary Information for "Characterizing the regulatory logic of transcriptional control at the DNA sequence level by ensembles of thermodynamic models"

[1]Departamento de Radiologia e Oncologia, Faculdade de Medicina, Universidade de Sao Paulo, Av. Dr. Arnaldo, 455 - Cerqueira Cesar, 01246 903, Sao Paulo, Brazil, [2]Instituto do Cancer do Estado de Sao Paulo Icesp, Hospital das Clinicas da Faculdade de Medicina da Universidade de Sao Paulo FMUSP HC, Av. Dr. Arnaldo, 251 - Cerqueira Cesar, 01246-000, Sao Paulo, Brazil, [3]Escola de Artes, Ciencias e Humanidades, Universidade de Sao Paulo, Rua Arlindo Béttio, 1000 - Ermelino Matarazzo, 03828-000, Sao Paulo, Brazil, [4]School of Life Science, Handong Global University, 37554, Gyeong-Buk, Republic of Korea, [6]Department of Ecology & Evolution, University of Chicago, 1101 E 57th St, 60637, Illinois, USA, [7]Department of Statistics, University of Chicago, 5747 S Ellis Ave, 60637, Illinois, USA and [8]Department of Molecular Genetics and Cell Biology, University of Chicago, 920 E. 58th St, 60637, Illinois, USA

*To whom correspondence should be addressed.

**Contact:**

**2 System specification** **7**

**3 Supplementary Figures** **8**

**References** **9**

### 1 The Transcription Model

Our transcription model, originally described in Reinitz et al. [2003], is structured in layers, each of which describes a regulatory mechanism. Since the new algorithms and results presented in Sections 3 and 4 make extensive use of those layers, understanding their biological interpretation is necessary. The inputs of these layers are DNA regulatory sequence in open chromatin and the intranuclear concentrations of TFs and their corresponding PWM. The output is the average transcription rate. Since the lifetime of mRNA is short compared to the timescale of changes in the average rate of its production, the observed level of mRNA is proportional to its average production rate. Because this quantitative mRNA observations come from many embryos [Janssens et al., 2005, 2006, Kosman et al., 1998, Pisarev et al., 2008, Poustelnikova et al., 2004, Surkova et al., 2008], we also have information about the variance of the pattern. We do not currently directly consider the effects of transcriptional bursting [Bothma et al., 2014, Fukaya et al., 2016, Garcia et al., 2013, Lammers et al., 2020, Pimmett et al., 2021]. Our transcription data is obtained by averaging mRNA levels over about 10 nuclei at each position on the anterior-posterior (A-P) axis. The expression of *eve* is a function of A-P and not dorsal-ventral position, hence this spatial average is equivalent to a temporal average over bursts. Our fixed tissue data exhibits bursting in some nuclei in the form of transcriptional spots [Kim et al., 2013, Figure 1].

#### 1.1 Identification of binding sites

The first layer of our model is the identification of binding sites from DNA sequence. TF binding sites are predicted by means of the PWM of each TF and the DNA regulatory sequence. For a given TF and each finite piece of the sequence, the PWM attributes a score. This can be interpreted as either the probability that a binding site exists or as an affine transformation of the Gibbs free energy ($\Delta G$) in a model in which $\Delta G$ for the whole binding site is the sum of $\Delta G$ for binding to each base. We take the latter interpretation, and so the score provides the binding affinity $K$ of the site by the relationship $K = \exp(\Delta G/RT)$, where $R$ is Boltzmann's constant and $T$ is the temperature in degrees Kelvin. The affine parameters for obtaining $\Delta G$ from the score are obtained as part of the fitting process [Barr and Reinitz, 2017, Supplementary Information]. The binding affinities are the output of this layer.

#### 1.2 Calculation of site occupancy

This second layer of our model calculates the fractional occupancy $f$ of each binding site, a property that will be very important in the material in Sections 3 and 4. $f$, a number between 0 and 1, denotes the probability that a binding site is occupied at any given instant, or equivalently, the proportion of the given binding sites that are occupied in a solution containing many molecules of the DNA containing the binding site and of the given TF at equilibrium. This is the precise reason why this class of models is called thermodynamic [Bertolino et al., 2016, Fakhouri et al.,

2010, He et al., 2010, Janssens et al., 2006, Kazemian et al., 2010, Kim et al., 2013, Martinez et al., 2014, Reinitz et al., 2003, Samee and Sinha, 2014, Sayal et al., 2016, Segal et al., 2008]. The calculation of $f$ for a single site is given by $f = \frac{K[TF]}{1+K[TF]}$, where $[TF]$ is the concentration of the given transcription factor, and $K$ is the binding affinity given above. The $f$s for a long segment of DNA with many binding sites is a straightforward but notationally complex generalization of the above formula which takes into account the fact that some sites overlap, and hence cannot be simultaneously occupied. We also consider the fact that sometimes the free energy of binding of a TF to two sites $\Delta G_{12}$ is greater than the sum of the free energy of binding to each site separetedly, so that $\Delta G_{12} = \Delta G_1 + \Delta G_2 + \Delta G_{\text{coop}}$. We incorporate cooperativity only for the TF Bicoid (Bcd), for which there is concrete experimental evidence [Kim et al., 2013, Lebrecht et al., 2005]. See Barr and Reinitz [2017] Supplementary Information for a technically precise description of this calculation.

It is important to note that the fractional occupancies $f$ described above are dependent on both DNA sequence and the local concentration of TFs, which vary in a complex manner along the A-P axis (see Fig. 1.**A**). Hence the $f$s vary by position, and this is what enables us to link transcription state to individual binding sites [Kim et al., 2013]. Our algorithm complexity for calculating the $f$s is linear to the length of the sequence, permitting us to calculate occupancies in the presence of very large number of binding sites [Barr and Reinitz, 2017, Supplementary Information]. In the cited work, 2920 binding sites were identified by taking a log odds threshold (PWM score) of zero, from which the calculation of $f$ requires considering every possible binding configuration. This calculation is rendered highly tractable by our algorithm, but the subsequent calculation of $f$ reveals a much smaller set of sites which actually influence transcription in an A-P axis position dependent manner. This establishes the need for a fine scale analysis of the regulatory profile within a gene locus and the importance of a sequence level model for transcriptional regulation

#### 1.3 The Roles of TFs

The model layers discussed below involve the physiological actions of the bound TFs as represented by $f$ at each binding site and at each position of the embryo. These TFs have differing physiological roles, which in the model described here can be derived from experiment, although it is feasible in other biological contexts to determine them directly by fitting to data [Bertolino et al., 2016]. The roles we consider here are activators, coactivators, and quenchers (short range repressors). In the biological context that we consider, transcription takes place because of activation. In the last layer of the model, the sum of bound activators $f$s will, after being modified by layers of regulation to be described below, feed into a DNA segment competition for interaction with the promoter, and an Arrhenius mechanism for a diffusion-limited enzyme, which we take to be RNA Polymerase II.

In what comes next we will describe layers of the model which begin from a picture of $f$s, to each of which is assigned a role of activator or quencher. In addition, activators may have an additional role as a coactivator. The phenomenological layers described below all involve modification of activators. In the third layer, coactivators convert sufficiently close quenchers into activators [Kim et al., 2013]. In the fourth layer, we calculate the effects of quenchers, which are repressors that have a limited range of action on the DNA, in nullifying the action of nearby activators, including those that have been subject to coactivation. All of these mechanisms phenomenologically represent the observed fact that multiple bound sites are required to repress or coactivate, that is to say that genetic function resides in multiple binding sites, the hallmark of a "billboard enhancer" [Kulkarni and Arnosti, 2005]. The fifth layer describes competition among enhancers to interact with the basal promoter. The model's ability to describe each site's physiological action individually will be essential in Section 3 and 4.

The TFs in the current application consist of the activators Bcd, Caudal (Cad), and *Drosophila*-STAT (Dst). Of these, both Cad and Bcd are coativators which act only on Hunchback (Hb). Kruppel (Kr), Giant (Gt), Knirps (Kni), Tailless (Tll), and Hb are quenchers.

#### 1.4 Coactivation

The layer of the model that calculates the effects of coactivation is a convolution over the fractional occupancies over the entire control region considered, $f_{i[m_i,n_i;a_i=\mathrm{Hb}]}$, where in this full notation $i$ is the numerical identifier of the site, which has 5' boundary at base $m_i$, 3' boundary at base $n_i$ and binds TF $a$, which in this case is Hb. These sites, before coactivation, all have a pure quenching activity. Coactivation is implemented by lowering the quenching activity of the physical $f_i$ subject to the constraint that

$$f_i = f_i^Q + f_i^A. \tag{1}$$

Thus, $f_i^Q$ is lowered by

$$f^Q_{i[m_i,n_i;a_i]} = f_{i[m_i,n_i;a_i]} \prod_k \left(1 - g(d_{ik}, D_c, 50) E^C_{a_k} f_{k[m_k,n_k;a_k]}\right), \tag{2}$$

where the product is taken over all $k$ sites in a control region that bind $\{\mathrm{Bcd}, \mathrm{Cad}\}$. The product over $k$ takes into account the fact that multiple sites are required for the action of coactivation in reducing quenching activity, since the contribution of a single coactivator may be close to unity, but the product of the activities of many coativators, each contributing a factor between 0 and 1 will be much closer to 0. Each coactivator $a$ has an efficiency $E^C_a$, determined by fitting over the range 0 to 1. The fact that coactivators act over a distance somewhat less than 250bp [Kim et al., 2013], is represented by $g(d_{ik}, D_c, 50)$. The function $g$ ranges between 0 and 1, and its arguments are as follows. $d_{ik}$ is the distance between the two closest borders of the binding sites $i$ and $k$ in basepairs. The shape of $g$ is trapezoidal and symmetric around site $i$. The second argument, $D_c$, is the distance from site $i$ where $g = 1$, and by symmetry the entire region where $g = 1$ is $2D_c$ long. The third argument, 50, gives the length of the linearly decreasing part of the trapezoid on both sides of site $i$. The experimentally observed constraint on $g$, and its uncertainty, are represented by allowing $D_c$ to vary during fitting within the range of 100bp and 200bp. Because of the constraint given in Eq.

(1), the activator activity of the bound TF is given by

$$f_i^A = f_i - f_i^Q. \tag{3}$$

#### 1.5 Quenching

This layer represents the effect of quenching (short range repression)[Fakhouri et al., 2010]. In Section 3 and 4 this level of the model describes the effects of repression on the $f^A$ defined up to this point. Short range repression is a key to understanding the importance of enhancers, since the range implies that a repressor in one enhancer does not have an effect on other enhancers. We note that short range repression acts on both dedicated and coactivated activators, but we do not allow the remaining $f^Q$ in a partially coactivated quencher to quench itself, thus keeping the graph of molecular interactions free of cycles. We denote an activated or coactivated $f$ that has been subject to quenching as an effective occupancy $F$. The effects of quenching are described by a convolution similar to that used to calculate coactivation, given by

$$F_i = f_i^A \prod_k \left(1 - g(d_{ik}, 100, 50) E_{a_k}^Q f_k^Q\right), \tag{4}$$

where the meanings of terms are very close to those in Eq. (1). $E_a^Q$ denotes the quenching efficiency of TF $a$, and is a number between 0 and 1 and hence determined by fits over the range 0 to 1. The convolution over $k$ has the same meaning as in Eq. (1), but there is more experimental information about the range of quenching [Fakhouri et al., 2010, Hewitt et al., 1999, Janssens et al., 2006], the second and third arguments of $g$ are experimentally determined and not fitted.

#### 1.6 Transcription adaptors recruiting

The effective occupancy of the DNA binding sites by activators $F$ provide the weights for computing their resulting capacity of recruiting the adaptors that will interact with the basal transcriptional machinery acting on the promoter. Let $E_{a_k}^A$ denote the adaptor attraction efficiency of the bound activator $a_k$. Then, the number of recruited adaptors to a sequence ranging from bp $p$ to $q$ becomes

$$N_{[p,q]} = \sum_k F_k E_{a_k}^A I(k, p, q), \tag{5}$$

where $I(k, p, q)$ indicates if site $k$ lies within $p$ and $q$:

$$I(k, p, q) = \begin{cases} 1, & m_k \geq p, n_k < q, \\ 0, & \text{otherwise.} \end{cases} \tag{6}$$

Previous formulations of the transcription model aimed at quantifying the effect of shorter DNA regions on promoter activity. Thus, it was assumed that all bound adapters affect the transcription rate [Janssens et al., 2006, Kim et al., 2013, Martinez et al., 2014, 2013, Reinitz et al., 2003]. Here, we apply the transcription model for predicting mRNA production rate as guided by the whole locus of *eve* gene. Hence, we consider that different DNA regions bound to adaptors compete for interacting with the basal transcriptional machinery [Barr and Reinitz, 2017].

#### 1.7 Enhancer competition

This layer of the model represents the fact that enhancers in general cannot all interact with the basal transcription complex at once. Indeed, repressors and coactivators have a limited range of action, and thus do not have activity which scales with the size of the DNA control region being modeled. The absence of a length action limit of controlling DNA would lead all activators to have an infinite range of action onto a promoter. As a consequence, larger control DNA segments would have greater activating power, and that does not correspond to experiment [Kim et al., 2013]. Finite range control DNA is modeled by the window size parameter $\alpha$. Additionally, there is ample evidence that only a single enhancer can interact with the basal promoter at once [Barr and Reinitz, 2017]. We represent this by weighting the probability of interaction of a particular segment of DNA with the basal promoter by the amount of adaptors acting on that segment. This mechanism, first presented in Barr and Reinitz [2017], is given by

$$T_{[m,m+\alpha]} = \frac{\beta N_{[m,m+\alpha]}}{1 + \sum_{n=1-\alpha}^{l} \beta N_{[n,n+\alpha]}}, \tag{7}$$

where $\beta$ is a parameter to be determined by optimization and $l$ is the whole locus length. This layer of the model represents the fact that the six enhancers of the *eve* locus individually regulate trancription at specific regions along the AP axis. The action of segment $[m, m+\alpha]$ has a relative duration $T_{[m,m+\alpha]}$ which depends on the amount of adaptors $N_{[m,m+\alpha]}$ that it recruits. The diffusion-limited Arhenius rate law will govern the effect of a segment on driving transcription.

#### 1.8 Diffusion-limited Arhenius rate law

A segment $[m, m+\alpha]$ interacting with the basal promoter induces mRNA synthesis at rate

$$R_{[m,m+\alpha]} = \frac{R_{\max}}{1 + \exp\left(\theta - N_{[m,m+\alpha]}\right)}, \tag{8}$$

where $R_{max}$ is the maximum value of transcription and $\theta$ is a positive constant denoting the energy barrier when no activator is bound to the DNA. Note that $N_{[m,m+\alpha]}$ also indicates the reduction of the energy barrier of transcriptional activation induced by the adaptors action and that $N_{[m,m+\alpha]} = 0$ enables one to set the basal transcription rate in terms of $\theta$. The scale of $N_{[m,m+\alpha]}$ and $\theta$ are set by optimization.

#### 1.9 Rate of transcription

The resulting rate of transcription in a position AP as induced by a whole locus of length $l$ is given by the sum of the action of each control DNA region balanced by its relative duration, which results in

$$R_{\text{Total}} = \sum_{m=1-\alpha}^{l} T_{[m,m+\alpha]} R_{[m,m+\alpha]}. \tag{9}$$

As previously discussed, the half-life of the transcripts is short in comparison with the timescales involved in transcription. Hence we set

$$[\text{mRNA}] \propto \frac{d}{dt}[\text{mRNA}] = R_{\text{Total}} \tag{10}$$

as the amount of mRNA's observed in a given AP position of the embryo.

The model presented here was validated experimentally in *D. melanogaster* embryos using *eve* as an experimental system [Barr and Reinitz, 2017, Janssens et al., 2006, Kim et al., 2013]. The multiple layers of the model discussed here enable the production of a complex pattern of expression based on the regulation of promoter activity by eight transcription factors. It provides the tools to aid on interpretation of experimental data aiming at identifying the major drivers of transcription in a position AP of the embryo. In the next subsection we summarize the biological implications of the transcription model.

### 2 System specification

Our system of five machines run Ubuntu 20.04 LTS Operational System. They have AMD Ryzen Threadripper CPUs, one have the model 3960X 24-Core Processor with 125Gb of RAM while the other four have the model 3990X 64-Core Processor with 252Gb of RAM. In our system, the $\sim 2000$ of serial simulated annealing took about 4 days to be completed in batches of 16 simultaneous runs by each machine.

### 3 Supplementary Figures

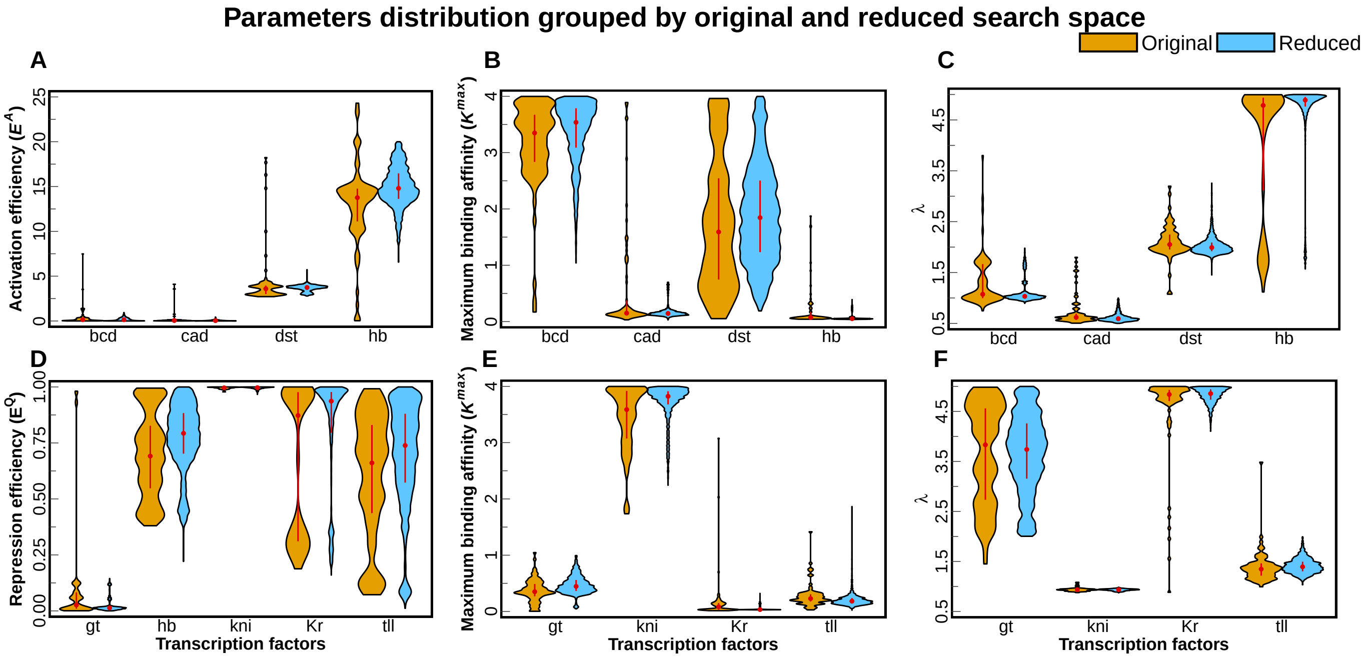

Supplementary Figure 1: Distribution of ensemble parameter values. The violin plots shows the efficiency, maximum binding affinity and $\lambda$ parameter values distribution for activators (A–C) and repressors (D–F). Orange (blue) violins for each TF, in x-axis, are the distribution values before (after) apply the reduction of the search space and a new round of SA. Red point inside the violins show the median while the straight line range between the first and third quartile.

### References

K. A. Barr and J. Reinitz. A sequence level model of an intact locus predicts the location and function of nonadditive enhancers. *PLoS One*, 12:e0180861, 2017. doi:10.1371/journal.pone.0180861. PMCID:PMC5513433.

E. Bertolino, J. Reinitz, and Manu. The analysis of novel distal Cebpa enhancers and silencers using a transcriptional model reveals the complex regulatory logic of hematopoietic lineage specification. *Developmental Biology*, 413:128–144, 2016. doi:10.1016/j.ydbio.2016.02.030. PMCID:PMC4878123.

J. P. Bothma, H. Garcia, E. Esposito, G. Schlissel, T. Gregor, and M. Levine. Dynamic regulation of *eve* stripe 2 expression reveals transcriptional bursts in living *drosophila* embryos. *Proceedings of the National Academy of Sciences USA*, 111:10598–10603, 2014. PMID:24994903 PMCID:PMC4115566 doi:10.1073/pnas.1410022111.

Walid D. Fakhouri, Ahmet Ay, Rupindar Sayal, Jacqueline Dresch, Evan Dayringer, and David N. Arnosti. Deciphering a transcriptional regulatory code: modeling short-range repression in the *Drosophila embryo*. *Molecular Systems Biology*, 6:341, 2010. PMID:20087339 PMCID:PMC2824527 doi:10.1038/msb.2009.97.

T. Fukaya, B. Lim, and M. Levine. Enhancer control of transcriptional bursting. *Cell*, 166:358–368, 2016. PMID:27293191 PMCID:PMC4970759 doi:10.1016/j.cell.2016.05.025.

Hernan G. Garcia, Mikhail Tikhonov, Albert Lin, and Thomas Gregor. Quantitative imaging of transcription in living *Drosophila* embryos links polymerase activity to patterning. *Current Biology*, 23:2140–2145, 2013. PMID:24139738 PMCID:PMC3828032 doi:10.1016/j.cub.2013.08.054.

X. He, M. A. H. Samee, C. Blatti, and S. Sinha. Thermodynamics-based models of transcriptional regulation by enhancers: The roles of synergistic activation, cooperative binding and short-range repression. *PLoS Computational Biology*, 6:e1000935, 2010. PMCID:PMC2940721.

G. F. Hewitt, B. Strunk, C. Margulies, T. Priputin, X. D. Wang, R. Amey, B. Pabst, D. Kosman, J. Reinitz, and D. N. Arnosti. Transcriptional repression by the *Drosophila* Giant protein: Cis element positioning provides an alternative means of interpreting an effector gradient. *Development*, 126:1201–1210, 1999.

H. Janssens, D. Kosman, C. E. Vanario-Alonso, J. Jaeger, M. Samsonova, and J. Reinitz. A high-throughput method for quantifying gene expression data from early *Drosophila* embryos. *Development, Genes and Evolution*, 215:374–381, 2005.

H. Janssens, S. Hou, J. Jaeger, A. R. Kim, E. Myasnikova, D. Sharp, and J. Reinitz. Quantitative and predictive model of transcriptional control of the *Drosophila melanogaster even skipped* gene. *Nature Genetics*, 38:1159–1165, 2006.

M. Kazemian, C. Blatti, A. Richards, M. McCutchan, N. Wakabayashi-Ito, A. S. Hammonds, S. E. Celniker, S. Kumar, S. A. Wolfe, M. H. Brodsky, and S. Sinha. Quantitative analysis of the *Drosophila* segmentation regulatory network using pattern generating potentials. *PLoS Biology*, 8:e1000456, 2010. PMCID:PMC2923081.

A. R. Kim, C. Martinez, J. Ionides, A. F. Ramos, M. Z. Ludwig, N. Ogawa, D. H. Sharp, and J. Reinitz. Rearrangements of 2.5 kilobases of noncoding DNA from the *Drosophila even-skipped* locus define predictive rules of genomic *cis*-regulatory logic. *PLoS Genetics*, 9:e1003243, 2013. PMCID:PMC3585115.

D. Kosman, S. Small, and J. Reinitz. Rapid preparation of a panel of polyclonal antibodies to *Drosophila* segmentation proteins. *Development, Genes and Evolution*, 208:290–294, 1998.

Meghana M. Kulkarni and David N. Arnosti. *cis*-Regulatory logic of short-range transcriptional repression in *Drosophila melanogaster*. *Molecular and Cellular Biology*, 25:3411–3420, 2005.

Nicholas C. Lammers, Vahe Galstyan, Armando Reimer, Sean A. Medin, Chris H. Wiggins, and Hernan G. Garcia. Multimodal transcriptional control of pattern formation in embryonic development. *Proceedings of the National Academy of Sciences*, 117:836–847, 2020.

D. Lebrecht, M. Foehr, E. Smith, F. J. P. Lopes, C. E. Vanario-Alonso, John Reinitz, D. S. Burz, and S. D. Hanes. Bicoid cooperative DNA binding is critical for embryonic patterning in *Drosophila*. *Proceedings of the National Academy of Sciences USA*, 102:13176–13181, 2005.

Carlos Martinez, Ah-Ram Kim, Joshua S. Rest, Michael Ludwig, Martin Kreitman, Kevin White, and John Reinitz. Ancestral resurrection of the *Drosophila* S2E enhancer reveals accessible evolutionary paths through compensatory change. *Molecular Biology and Evolution*, 31:903–916, 2014. doi: doi:10.1093/molbev/msu042. PMCID:PMC3969564.

Carlos A. Martinez, Kenneth A. Barr, Ah-Ram Kim, and John Reinitz. A synthetic biology approach to the development of transcriptional regulatory models and custom enhancer design. *Methods*, 62:91–98, 2013. doi: doi:10.1016/j.ymeth.2013.05.014. PMCID:PMC3924567.

Virginia L Pimmett, Matthieu Dejean, Carola Fernandez, Antonio Trullo, Edouard Bertrand, Ovidiu Radulescu, and Mounia Lagha. Quantitative imaging of transcription in living Drosophila embryos reveals the impact of core promoter motifs on promoter state dynamics. *Nature communications*, 12:4504, 2021.

A. Pisarev, E. Poustelnikova, M. Samsonova, and J. Reinitz. FlyEx, the quantitative atlas on segmentation gene expression at cellular resolution. *Nucleic Acids Research*, 37:D560–D566, 2008. PMCID:PMC2686593.

E. Poustelnikova, A. Pisarev, M. Blagov, M. Samsonova, and J. Reinitz. A database for management of gene expression data in situ. *Bioinformatics*, 20:2212–2221, 2004.

J. Reinitz, S. Hou, and D. H. Sharp. Transcriptional control in *Drosophila*. *ComPlexUs*, 1:54–64, 2003.

M. A. H. Samee and S. Sinha. Quantitative modeling of a gene's expression from its intergenic sequence. *PLoS Computational Biology*, 10:1–21, 2014.

R. Sayal, J. M. Dresch, I. Pushel, B. R. Taylor, and D. Arnosti. Quantitative perturbation-based analysis of gene expression predicts enhancer activity in early *Drosophila* embryo. *eLife*, 5:e08445, 2016. PMID:27152947 PMCID:PMC4859806 doi:10.7554/eLife.08445.

E. Segal, T. Raveh-Sadka, M. Schroeder, U. Unnerstall, and U. Gaul. Predicting expression patterns from regulatory sequence in *Drosophila* segmentation. *Nature*, 451:535–540, 2008. PMID:18172436 doi:10.1038/nature06496.

S. Surkova, D. Kosman, K. Kozlov, Manu, E. Myasnikova, A. Samsonova, A. Spirov, C. E. Vanario-Alonso, M. Samsonova, and J. Reinitz. Characterization of the *Drosophila* segment determination morphome. *Developmental Biology*, 313(2):844–862, 2008. PMCID:PMC2254320.
